# Interface residues that drive allosteric transitions also control the assembly of L-lactate Dehydrogenase

**DOI:** 10.1101/361550

**Authors:** Jie Chen, D. Thirumalai

## Abstract

The allosteric enzyme, L-lactate dehydrogenase (LDH), is activated by fructose 1,6-metaphosphate (FBP) to reduce pyruvate to lactate. The molecular details of the FBP-driven transition between the low affinity T-state to the high affinity R-state in LDH, a tetramer composed of identical subunits, are not known. The dynamics of theT→R allosteric transition, investigated using Brownian dynamics (BD) simulations of the Self-Organized Polymer (SOP) model, revealed that coordinated rotations of the subunits drive the T→R transition. We used the structural perturbation method (SPM), which requires only the static structure, to identify the allostery wiring diagram (AWD), a network of residues that transmits signals across the tetramer, as LDH undergoes the T→R transition. Interestingly, the residues that play a major role in the dynamics, which are predominantly localized at the interfaces, coincide with the AWD identified using the SPM. The conformational changes in the T→R transition start from the region near the active site, comprising of helix *α*C, helix *α*1/2G, helix *α*3G and helix *α*2F, and proceed to other structural units, thus completing the global motion. Brownian dynamics simulations of the tetramer assembly, triggered by a temperature quench from the fully disrupted conformations, show that the bottleneck for assembly is the formation of the correct orientation between the subunits, requiring contacts between the interface residues. Surprisingly, these residues are part of the AWD, which was identified using the SPM. Taken together, our results show that LDH, and perhaps other multi-domain proteins, may have evolved to stabilize distinct states of allosteric enzymes using precisely the same AWD that also controls the functionally relevant allosteric transitions.

## Introduction

Allosteric transitions, resulting in signal transmission in proteins and their complexes in response to external perturbations, such as ligand binding or mechanical force, is pervasive in biology^1–3^. The scales over such signaling occurs could vary from a few nanometers (hemoglobin for example) to several nanometers (dynein). This in itself is interesting, if not surprising, because these length scales are typically larger than the size of the individual subdomains, which means that allosteric signals are transmitted across interfaces in multi-domain proteins. Many of the multi-domain proteins undergo large scale conformational fluctuations in response to local perturbations, driving the system from one state to another during a particular reaction cycle that is typically associated with their functions. Because of the pervasive nature of signal transmission in these mesoscale systems, it stands to reason that there must be general molecular principles governing allostery, as was illustrated beautifully, using essentially symmetry arguments, in the celebrated Monod-Wyman-Changeux (MWC) theory^4^. The ubiquitous observation of allosteric transitions from molecules, with simple folds to complex multi-domain proteins, has given rise to a large enterprise in search for molecular explanation of allostery using theory^5–7^, experiments^8–10^, and computer simulations^11,12^.

Here, we use computational techniques to probe the dynamics of allosteric transitions in L-lactate dehydrogenase (LDH), an important enzyme^13,14^, but one which has received surprisingly little attention from the biophysical community. The enzyme LDH catalyzes the interconversion of pyruvate and lactate^15^. Some bacterial LDH are allosteric enzymes activated by fructose 1,6-bisphosphate (FBP)^16–18^. In contrast to non-allosteric vertebrate LDHs, allosteric LDHs assemble as tetramers^15,18–20^ composed of identical subunits related through three twofold axes labeled P and Q (Fig. 1) and R, following the Rossmann convention ^15^.

**Figure 1:**
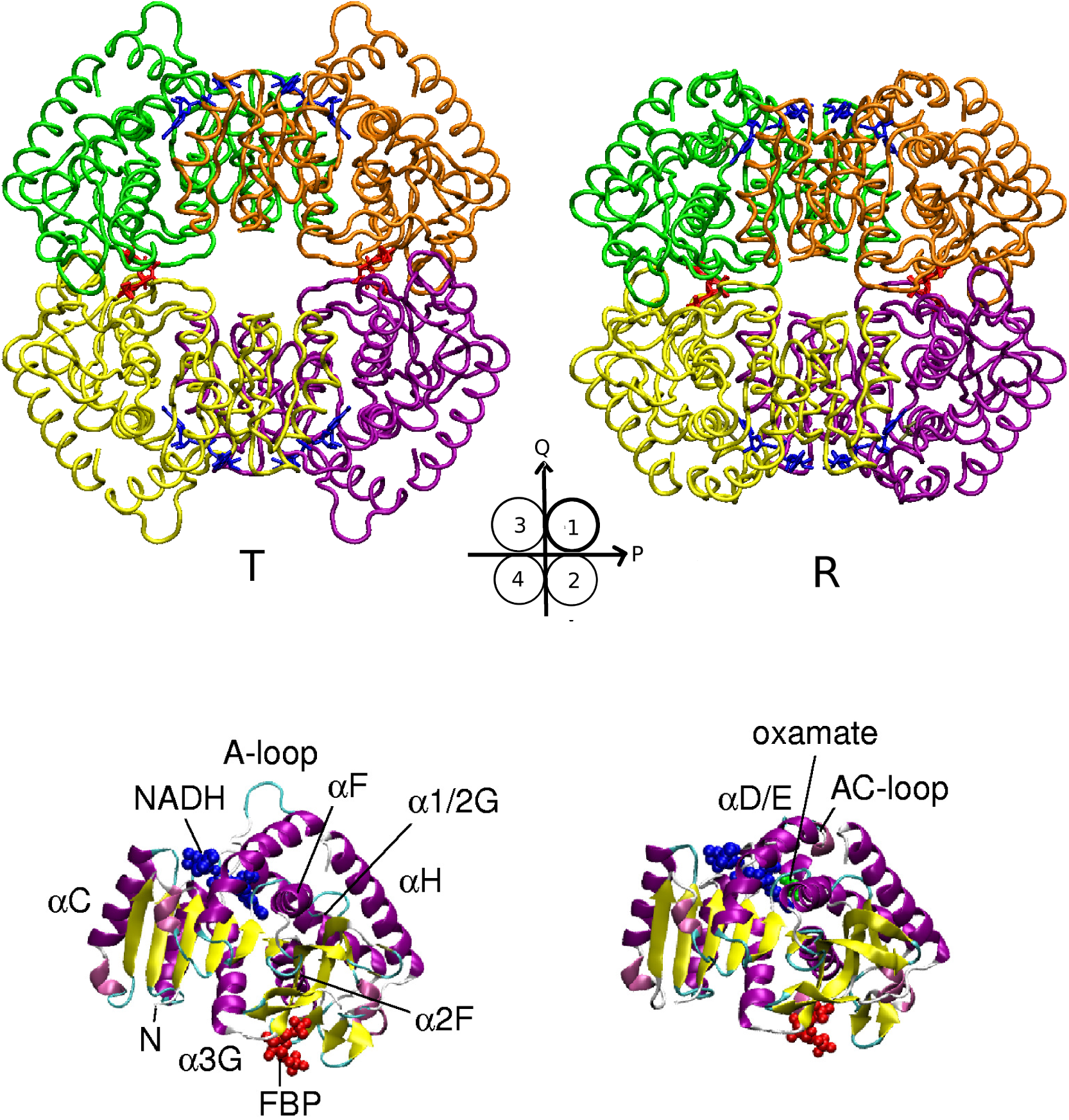
Structural representation of the T (left) and R (right) states go tetrameric LDH. The structures are colored for different chains (the organization of the four subunits is shown schematically in the middle in circles) with bound ligands in licorice (red for FBP and blue for NADH). In the lower part are the cartoon representations of the subunits of the T and R states. Key secondary structural elements are labeled. Ligands NADH and FBP are displayed using blue and red spheres.

LDH from *Bifidobacterium longum* (BLLDH) is an allosteric enzyme whose crystal structures in the low (T-state) and high (R-state) affinity states have been determined^16^ (Fig. 1). The FBP binding sites consist of four positively charged residues, R173 and H188 from two P-axis-related subunits. Upon FBP binding, the charge repulsion between R173 and H188 from the two P-axis-related subunits is diminished. As a result, the open conformation of LDH in the T-state changes to the closed conformation in the R-state. Comparison of the crystal structures of the T- and R-states of LDH shows that the T→R transition involves two successive rotations of the subunits: first rotation is ≈ 3.8° about an axis close to H188, caused by binding of FBP on the P-axis-related subunit interface. The second is a ≈ 5.8 ° rotation about the P-axis, which creates changes in the hydrophobic interactions on the Q-axis related subunit interface to which the substrate would bind. These quaternary structural rearrangements reflect the allosteric control of LDH, triggered by switching between two distinct states. The mechanism underlying this transition, which is assumed to preserve the symmetry of the allosteric states, is in accord with the MWC model^4,21^ in two important ways. First, regulatory proteins are oligomers formed by identical subunits with underlying symmetry. Second, the interconversion of oligomers between discrete conformational states is independent of the presence of ligands, which is the basis of conformational selection that has drawn considerable attention in recent years. Of course, in the presence of ligands the transition occurs more readily leading to enhanced stability of the R state.

Changes in the tertiary structure and cooperative interactions are also amplified to buffer the quaternary structural changes. By comparing the known crystal structures of LDH, the conformational changes of the structural units are found in the sliding of helix *α*C on the Q-axis-related subunits interface, causing helix *α*1/2G to kink leading to the shift in the carboxy terminal out of the active site (see Fig. 1). Therefore, *α*C sliding is thought to control the affinity of the substrate^16^. However, there is no information about the dynamics of helix sliding nor about the conformational changes that propel the T→ transition.

Although some insights into allostery in LDH have been gleaned from the available structures, the dynamics of the T→R transition in LDH has not been examined. Here, we use several methods to produce a comprehensive picture of the T→R transition. (i) First, we used the Structural Perturbation Method (SPM)^22,23^ to show that the allosteric communication pathways in the T→R transition involves a network of key residues, referred to as the Allosteric Wiring Diagram (AWD), located at the subunit interface. The SPM utilizes only the static structures and predicts the response to local perturbation, which is calculated based on a normal mode analysis. (ii) Sometime ago, we introduced a coarse-grained Self-Organized Polymer (SOP) model^24–27^, which has been used to describe the dynamical transitions between distinct allosteric states in a variety of multi subunit systems^25–27^. These simulations, which predict the dynamics of key events driving the transition between two states, revealed breakage of dynamical symmetry of the seven subunits during the allosteric transitions in the bacterial chaperonin GroEL, triggered by ATP binding and hydrolysis^26^. Here, we use similar techniques to probe allosteric transitions in LDH. The conformational changes monitored using a number of quantitative measures, reveal that the key events in the T→R transition involve rotations of the subunits, which drive the needed tertiary structure changes. The allosteric communication pathways in the T→R transition involving a network of key residues located at the subunit interface, calculated from the dynamical simulations coincide with the AWD residues determined using the SPM^22,23^ predictions. (iii) We also investigated the assembly of LDH starting from disordered high temperature structures by a quench to low temperatures. Although there are parallel assembly routes, in the dominant path the subunits fold first and subsequently they reorient themselves to achieve the correct orientational registry. Remarkably, the residues in the AWD not only drive the allosteric T→R transition but are also involved in the assembly of LDH monomers. Taken together our results suggest that the dominant residues that control oligomer formation and allosteric transitions may have co-evolved to simultaneously optimize dynamics and assemblies of multi-domain proteins.

## Methods

### Allosteric Wiring Diagram (AWD) from the Structural Perturbation Method (SPM)

In order to identify the key residues in the allosteric transition, we used the SPM, which begins with an elastic network representation of the states^22,23,28^. We used a harmonic potential with a single force constant for the pairwise interaction between all *C_α_* atoms that are within a cutoff distance (*R_c_* = 10Å) to build an elastic network^29–31^. We note parenthetically that related ideas have been proposed more recently^32,33^.The elastic energy of the network is,

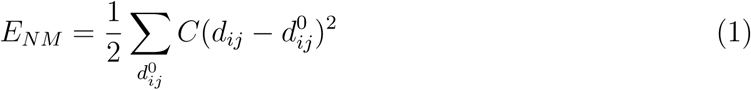

where
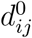
is the distance between residue i and residue j in the LDH native structure, and *C* is a spring constant, whose value is chosen to match the crystallographic B-factors in a given allosteric state.

We performed standard normal mode analysis using Eq. (1) to get the eigenvectors of the 30 lowest modes, which could represent the function-related motions^29–31^. Using, *v_ij_*, the eigenvector of mode *j*, we computed the overlap function describing the conformational changes,
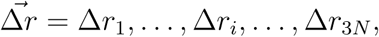
from the T to R state using^22,34,35^,

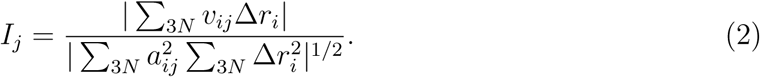

We identify the dominant mode connecting the T→R transition as the one with the largest value of *I_j_*.

To determine the AWD^22,23^, we assessed the effect of a point mutation on the residue at position *i* on T→R transition by adding a perturbation to *E_network_* given by,

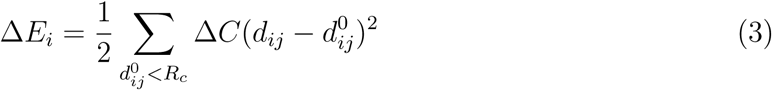

where *j* is the index of the residue that has contacts with residue *i* and Δ*C* = − *C*/2. The response of the mode M, with the largest overlap, to the point mutation is calculated using
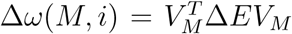
(T is the transpose of the matrix with elements *v_ij_*). The residues that carry the largest values of the stored elastic energy (Δ*ω*(*M*, *i*)) are considered to be part of the AWD. The details of the theory are outlined elsewhere^23,36^.

### Self-organized polymer (SOP) model for allosteric transitions

The long-time scales in the ligand-driven transitions make it impossible to carry out meaningful computations using atomically detailed simulations, thus making it is necessary to use a coarse-grained models^37,38^. We use the self-organized polymer (SOP) model^24^, which has been remarkably successful in applications to protein and RNA folding as well in probing allosteric transitions in molecular machines^27,39^, for probing the T→R transition in LDH. In the simplest version of the SOP model, the structure of the complex is represented using the *C_α_* coordinates. The allosteric state-dependent energy function in the SOP representation, in terms of the *C_α_* coordinates *r_i_*(*i* = 1, 2, …*N*), with *N* being the number of amino acids, is

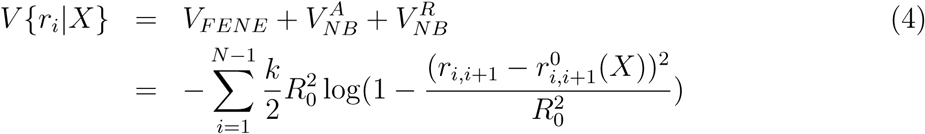

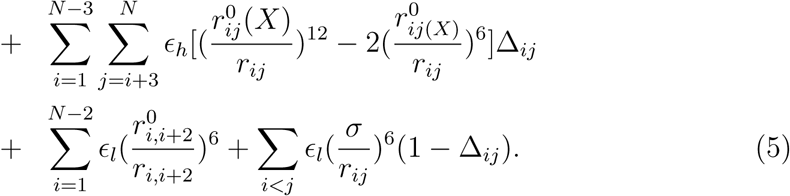

The label *X* in the above equation, refers to the allosteric state, T or R. In Eq. (4), *r*_*i*, *i*+1_ is the distance between two adjacent *C_α_*-atoms and *r*_*i*,*j*_ is the distance between the *i^th^* and *j^th^ α*-carbon atoms. The superscript 0 denotes their values in the allosteric state, *X*. The first term in Eq. (4), the finite extensible non-linear elastic (FENE) potential, accounts for chain connectivity. The stability of the state *X* is accounted for by the non-bonded interactions (second term in Eq.(4)), which are taken into account by assigning attractive interactions between residues that are in contact in *X*. The value of Δ_*i*,*j*_ is unity if the sites *i* and *j* are in native contact, and is zero otherwise. We assume that native contact exists if the distance between the *i^th^* and *j^th^ C_α_* atoms is less than 8Å. Non-bonded interactions between residues that are not in contact in *X* are repulsive (third term in Eq. (4)). The spring constant *k* in the FENE potential for stretching the covalent bond is 20 kcal/(mol · Å)^2^, and the value of *R*_0_, which gives the allowed extension of the covalent bond, is 2Å. These are the standard values used previously in several applications of the SOP model to allosteric transitions in a variety of systems^26,27,40^.

### Allosteric transition from the T→R state

In order to investigate the molecular basis of the T→R transition dynamics, we adopted a method previously used to probe allosteric transitions in the GroEL reaction cycle and myosin motors^26,27,40^. We assume that the local strain that LDH experiences when a ligand binds propagates faster across the structure than the time scales for conformational transitions leading to the R state from the T state. To observe transitions from one allosteric state (say T) to another (R for example), we start from a conformation corresponding to the T state. The transition is induced by using the forces calculated from the R the state as a derivative of the R state Hamiltonian, *V*{*r_i_*|*R*} (see Eq. (1)). The explicit equations of motion for the T→R transition are,

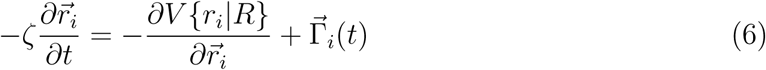

where *r_ij_* and
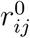
are the distances between residues *i* and *j* in a given conformation generated during the dynamics and the value in the allosteric state, respectively.

The T-state and R-state tetrameric structures of LDH are obtained by performing a symmetry operation for the monomeric LDH (PDB: 1LTH). We initiated Brownian dynamics simulations by first equilibrating the T-state LDH for 80 *μ*s. Subsequently, the equations of motion are integrated using Eq.6 for an additional 80 *μ*s. The integration step is h=0.16, except in the initial stages of the T→R transition during which it is changed to h=0.016 for 8 *μ*s to ensure that there are no numerical instabilities. All the simulations are done at *T* = 300*k*. Additional details of the simulations are described elsewhere^26^.

## Results

In order to probe the global motion of LDH during the T→R transition, we simulated the entire tetramer using Brownian dynamics, as described in the Methods section. The dynamics of the T→R transition is monitored by analyzing the time evolution of various distances and angles describing the LDH structure. The residue numbers in the following analysis are taken from the Protein Data Bank (PDB) where the convention for LDH numbering^15^ is given in parentheses.

### Tetramer dynamics decomposed into rotations about the PQ and QR planes

Binding of FBP ligand to LDH triggers quaternary conformational changes of LDH^16^. In order to probe the global motion of LDH during the allosteric T→R transition, we monitored the time-dependent changes in three angles *α*, *β*, and *γ* (see the legend to Fig. 2 for definitions), measuring the relative orientations of the subunits. The angle *α* is almost constant during the T→R transition, showing that the relative orientation of the two boundary helices in one subunit remains unchanged. The *β* angle on the other hand decreases by about 10 ° during the T→R transition, indicating a counter-clockwise rotation of subunit 1 and a clockwise rotation of subunit 2 (P-axis-related to subunit 1). In the RQ plane, the angle between helix *α2F* and R axis decreases from 23 to 14 °, indicating counter-clockwise rotations of subunits 1 and 2 around the RQ plane.

**Figure 2:**
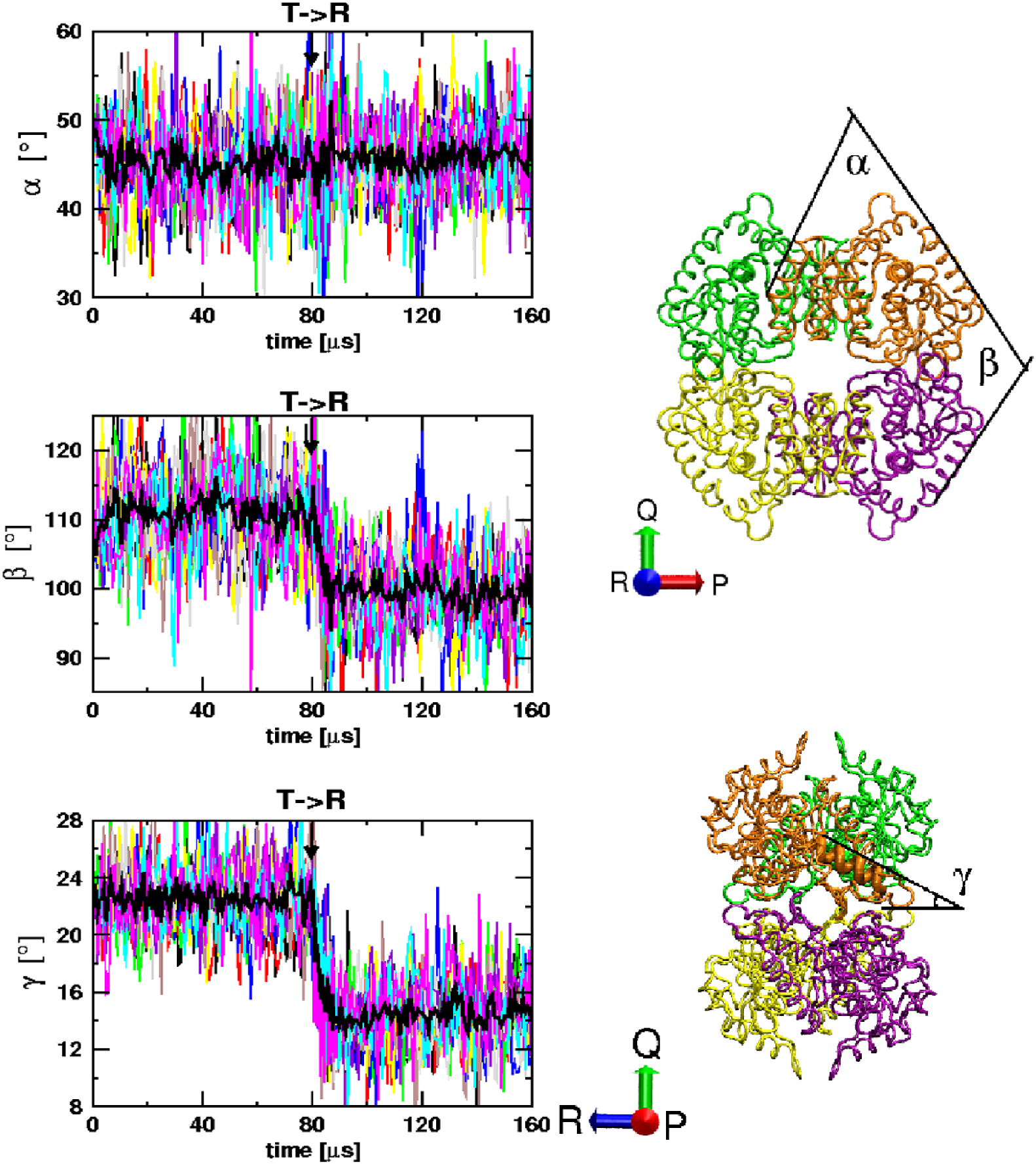
LDH dynamics monitored using the angles *α*, *β* and *γ*. An angle, labeled *θ*(*t*), is defined by
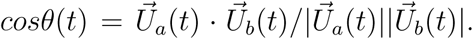
For angles *α* and *β*,
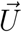
is defined as the projection of
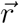
onto the plane perpendicular to axis *ê_R_* using
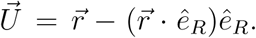
 For angle *α*,
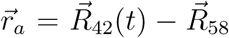
is the direction of helix *α*C, and
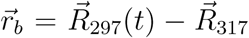
is the direction of *α*H. For angle
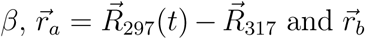
is same as
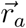
except it is taken from the P-axis related subunit. For angle *γ*, the
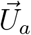
is defined as the projections of
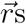
onto the plane perpendicular to axis *ê_P_* and
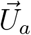
is axis *ê_R_*. The time-dependencies of *α*, *β* and *γ* for ten trajectories are plotted in different colors, with the black lines being the average over ten trajectories. Angles *α*, *β* and *γ* are indicated in the T state LDH on the right.

The combined results for the rotational changes Fig. 2 and the root mean square deviations (RMSDs) for the tetramer (Fig. 3A) and monomer (Fig. 3B) show that the overall conformational changes of the tetramer is more significant than the changes that occur in the individual subunits. Thus, the T→R transition in LDH may be viewed as a symmetry-preserving rigid body rotation with little, if any, partial unraveling of the monomers unlike in GroEL^26^. The global T→R transition is consistent with two-state kinetics, which follows from the observation that the kinetics of ensemble averaged RMSDs for tetrameric LDH can be fit by a single exponential function (Fig. 3C).

**Figure 3:**
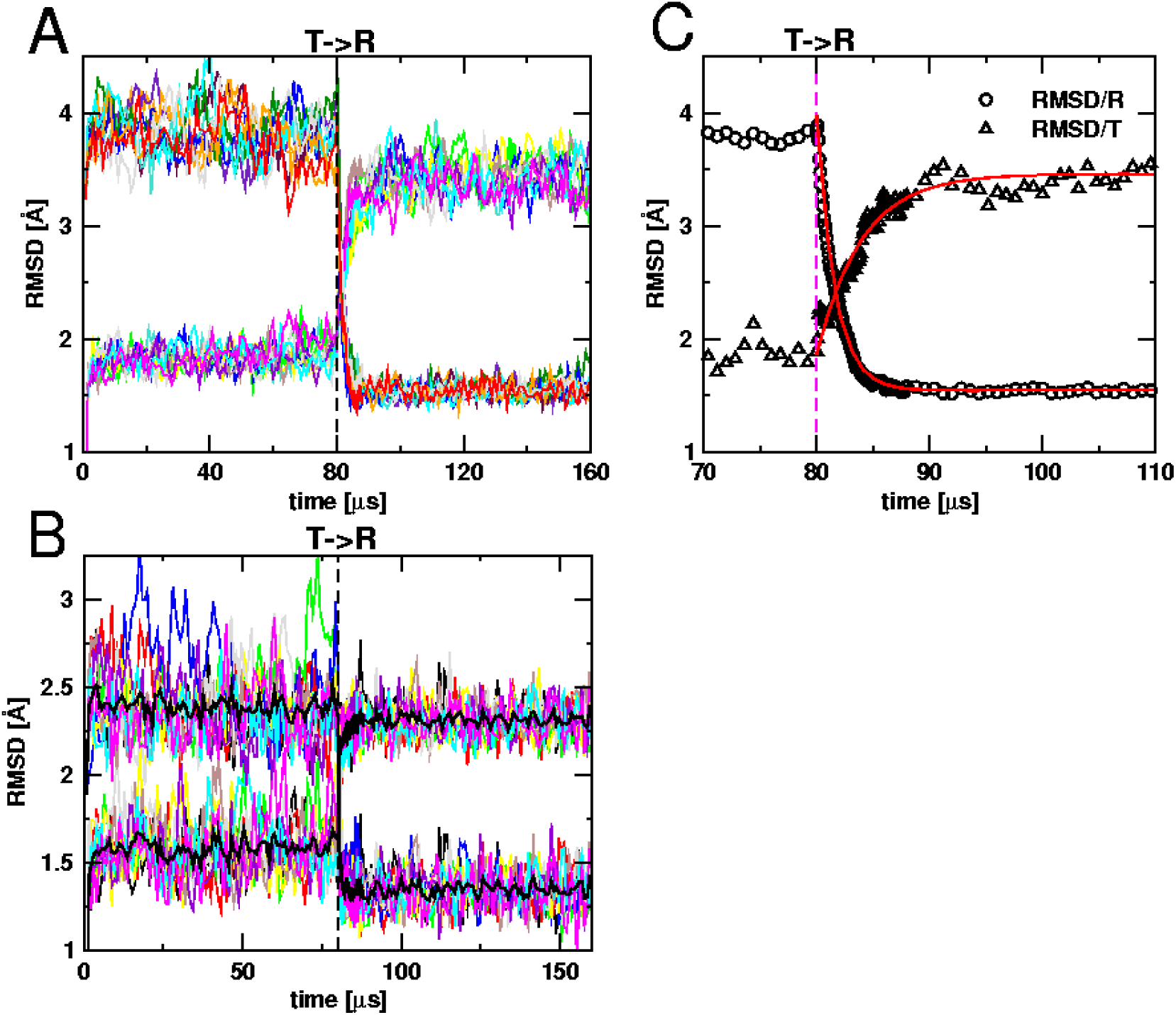
RMSD as a function of time. A: Tetramer RMSDs between the T and R states are plotted for ten trajectories in different colors. The transition to the R state is initiated at *t* = 80*μ*s. B: Time-dependent RMSDs for subunits of LDH for ten trajectories are plotted in colors with the black lines being the averages. C: Average of tetramer RMSDs obtained from ten trajectories. RMSD/T (with respect to the T state) is given in triangles and RMSD/R (with respect to the R state) is in circles. The two RMSD lines, fit to single exponential function, are shown as solid red lines. The fits are: < *RMSD/T*[] > = 3.46 − 1.60*e*^−*t*/3.95*μS*^) and < *RMSD/R*[] > = 1.55 + 2.44*e*^−*t*/1.60*μs*^.

### Fast and slow transitions associated with changes in the secondary structural elements

To elucidate the changes in the secondary structural elements (SSEs) that take place in the T→R transition, we compared the fate of the secondary structural elements at the transition state (TS). Such an analysis allows us to dissect the order in which these transitions occur in the T→R transition. Using the assumption that TS location is reached when *δ*^‡^ = |*RMSD/T*(*t_TS_*) − *RMSD/R*(*t_TS_*)| < *r_c_*, where *r_c_* = 0.2 and *t_TS_* is the time at which *δ*^‡^ < *r_c_*, we categorized the transition states of the eight helices in LDH and three loops (active loop (A-loop),active control loop (AC-loop) and flexible surface loop (FS-loop)), into two groups - those that undergo fast transitions, and the ones that change on longer time scales (Fig. 5). The categorization is predicated on the values of *t_TS_*. Helices *α*1/2G, *α*2F, *α*C, *α*3G, A-loop and AC-loop fall into fast transition category with *t_TS_* < 82*μs*. The other four helices and the FS-loop fall in the slow transition category. We note that all the SSEs involved the in the neighborhood of the active site undergo fast conformational changes during the T→R transition, which agrees with the intuition that binding of activators occurs rapidly in catalytic reactions. Our simulations show that the the order of allosteric transitions associated with SSEs is, {*α*1/2G, *α*2F, A-loop, *α*C, *α*3G, AC-loop, *α*D/E, *α*B, *α*H, FS-loop}, implying that there is a hierarchy in the internal dynamics governing the T→R transition, much like in Myosin V^40^.

### Allosteric signal transduction mechanism in LDH

To answer the question of how allosteric signal transmission occurs from the effector site to the active site, we probed the dynamics of the charge interactions on the P-interface around the FBP binding sites, the dynamics of the hydrophobic interactions on the Q-interface around the active sites, and the interactions inside a single subunit. The changes in interactions are monitored using the distances between two interacting residues, *r*_*i*,*j*_(*t*). We fit *r*_*i*,*j*_(*t* > *t_eq_*), where *t_eq_* is the time to equilibrate the tetramer, to single exponential functions, *r*_*i*,*j*_(*t*) = *a* + *bexp*(−*t/τ*). The characteristic time *τ* is a measure of how fast the interaction energy changes during the T→R transition. With this analysis method it is straightforward to identify the sequence of the interaction changes by comparing the *τ* values, thus providing quantitative insights into the allosteric signal transduction mechanism in LDH triggered by binding of FBP.

Our simulations show that at the Q-interface the distance changes are well fit using a sum of two exponential terms with a fast phase (*τ* ~ 0.1 − 0.3*μs*) and a slow phase *τ* ~ 1 − 5*μs*. In the fast phase, the hydrophobic interactions are modified before the interactions change at the active site. The slow phase lasts until the charge repulsion is neutralized by FBP on the P-interface. In detail, the hydrophobic interactions on the Q-interface between *α*C(Q) and the active control loop, *α*C(Q) and *α*2F change in the fast phase (see top and middle panels in Fig. 4), which causes contact formation between Arg171-Tyr190 and Asn140-His195 in the active site. Subsequently, the active control loop changes conformation resulting in a contact between Asp168-His195 forms, and then Ile240 moves out of the substrate binding pocket. These events results in the closing of the active loop.

**Figure 4:**
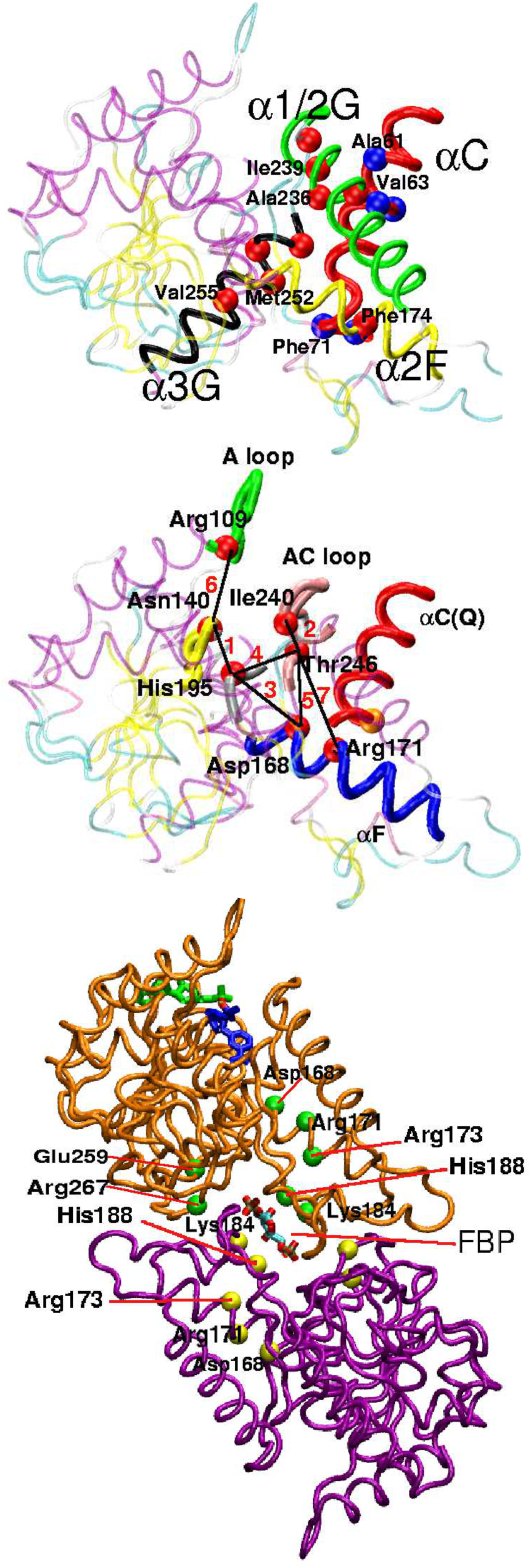
Changes in the interface hydrophobic interactions at the Q-interface in the T→R transition. (Top) A snapshot from the simulations shows that helix *α*C belonging to subunit 3 is in red with important hydrophobic residues displayed in blue spheres. Helix *α*2F, *α*1/2G and *α*3G of subunit 1 are shown in yellow, green and black, respectively. The relevant residues in these helices are shown in blue spheres and labeled. Other structural elements units except the three helices are colored according to their secondary structures and are shown in transparent for clarity. (Middle) Formation and rupture of contacts at the active site inside subunit 1 during the T→R transition. A few key residues are labeled and the sequence of changes of interactions are indicated with a number; 1 indicates the first event and 7, the last event. The numbering of the residues follows the conventional LDH numbering system. (Bottom) Changes in the electrostatic interactions on the P-interface in T→R transition. subunits 1 and 2 are shown in orange and purple with important charged residues shown using green and yellow spheres. All the residues are labeled and FBP is explicitly shown.

**Figure 5:**
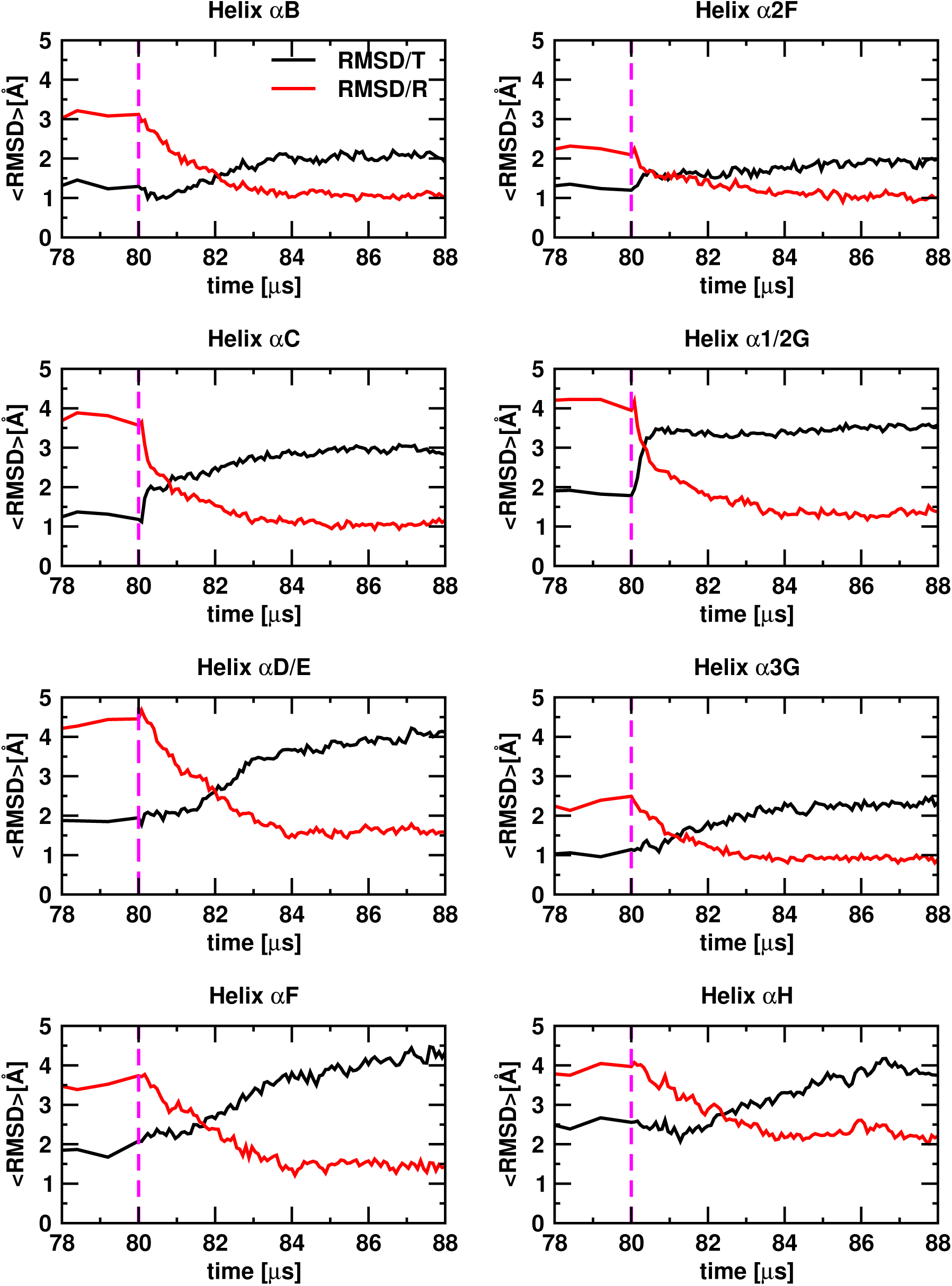
Averaged RMSD as a function of time for each helix. < *RMSD/T* > and < *RMSD/R* > are drawn in black and red lines, respectively. The time *t_TS_* is defined as the time where the two lines of RMSD cross (see the text). According to the values of *t_TS_* the eight helices are categorized into fast transition group (*t_TS_* < 82*μs*), and slow transition group (*t_TS_* < 82*μs*).

In the slow phase interactions on the Q-interface are altered. Electrostatic repulsion between residues Arg173 and His188 is reduced by binding of FBP, which leads to proximity of the two P-interface related subunits (see middle panel in Fig. 4). It is surprising that the distances between charged residues Asp168-Lys184, Arg170-Lys184 and Lys184-Lys184 do not increase monotonically. Instead, they increase rapidly to a value that is almost double their target distances in the R state, and then decrease. The increase occurs in the fast phase during which interaction changes on Q-interface change while the decrease in the distances in the slow phase indicates the stabilization of P-interface by FBP. These results show that *α*C(Q) slides along the Q-interface and changes the active area conformation that is stabilized by binding of FBP to the P-interface.

### Allostery Wiring Diagram and the structural origin of the slow transitions identified by the SPM

The immediate question that raises from the identified order of events discussed above is: what is the significance of the structural units whose transitions are slow? To answer this question, we performed NMA for a LDH tetramer to obtain the thirty lowest modes. For a large number of proteins it has bee shown that only a few low frequency modes are functionally relevant^22,28,41,42^. We found that for mode seven the overlap value is 0.75, implying that it contributes most to the conformational changes in the T→R transition (top panel in Fig. 6). Focusing on mode 7, we did site mutation for each residue of subunit 1 (Methods) in order to determine the AWD connecting the T and R states using the SPM equation (Eq. 3). The effect of mutation is evaluated by calculating the residue-dependent elastic energy change associated with mode 7 under perturbation (see Fig. 6). We identify seven peaks in Fig. 6, which are associated with the most important residues in the AWD. Interestingly, all these residues are around helix *α*D/E, *α*F, and *α*H, which are the structural elements (see Fig. 6) that undergo slow transitions. We surmise that these residues drive global motions in the allosteric transition.

**Figure 6:**
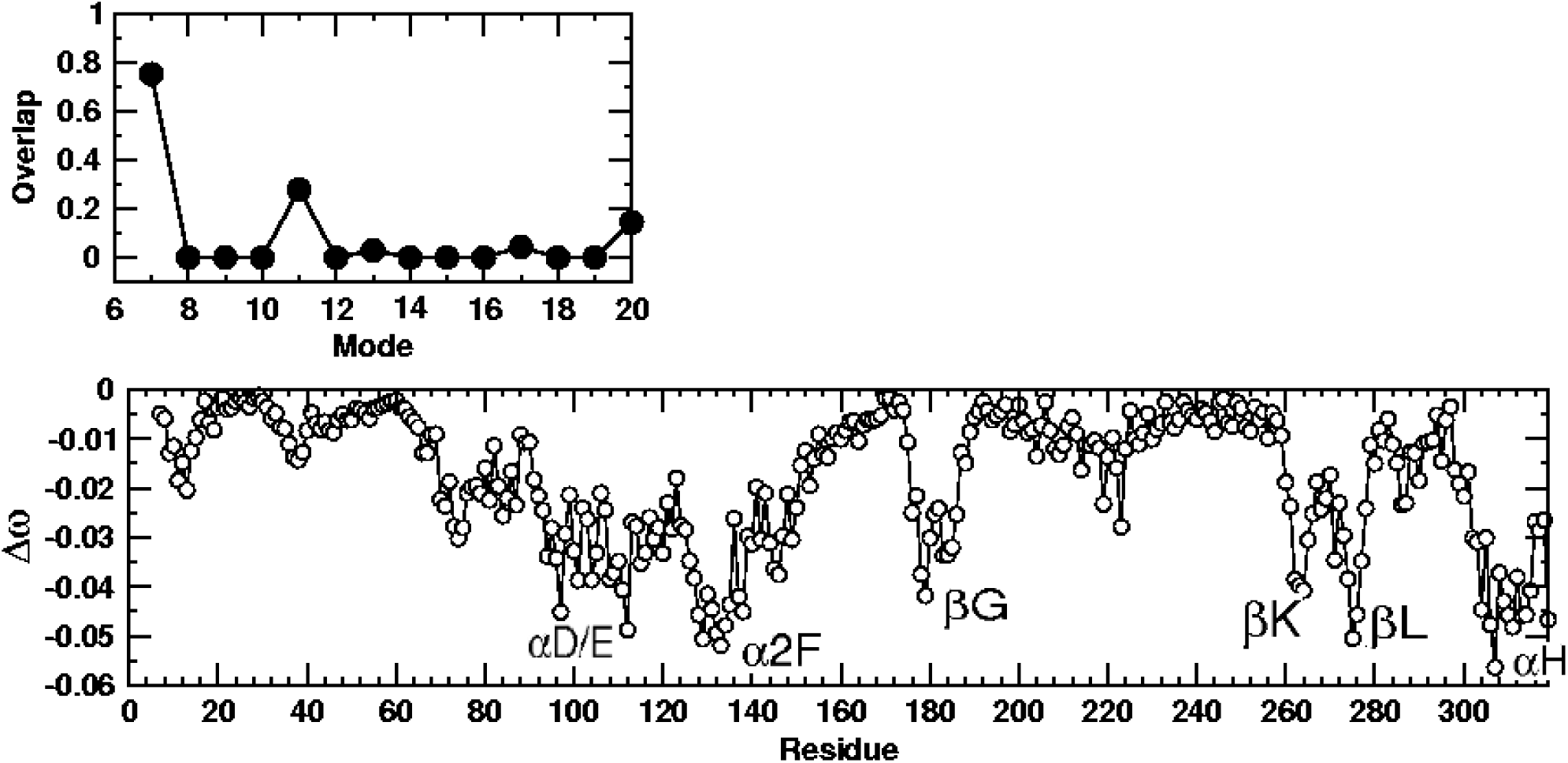
Allostery wiring diagram of LDH inferred from SPM. The overlaps of 20 lowest mode with the conformational change in the T→R transition are shown on top. Mode 7 has the highest overlap value, 0.75. In the bottom panel, the residue dependent stored elastic energy is plotted in response to perturbation is plotted for mode 7. Residues with large negative frequency changes are the most significant part of the AWD. Their locations are associated with secondary structural elements (helix *α*D/E, helix *α*2F, helix *α*H), belonging to the slow transition group.

### Rate determining step in the assembly is the orientational registry across the interface

In an attempt to further elucidate if the interactions of LDH across the interfaces also control assembly, we simulated the assembly of a LDH dimer containing Q-interface related subunits 1 and 3 in the R state. We first equilibrated the dimer for *t* < *t*_0_ then increased the temperature from 300K to 1200K in ten steps in the time interval, < *t*_0_ < *t* < *t*_1_. In the interval, *t*_1_ < *t* < **t**_2_, we decreased the temperature back to 300K to initiate the assembly of LDH. subsequently, we equilibrated the dimer in the duration *t*_2_ < *t* < *t*_3_. We set to = 50*μs*, *t*_1_ = 150*μs*, *t*_2_ = 250*μs* and *t*_3_ = 350*μs*. In order to keep the dimer intact, we included a weak FENE potential to constrain the centers of mass of the two subunits ^43^.

We monitored the assembly process using the fraction of contacts,
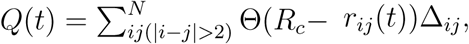
where *R_c_* is the cut-off distance for native contacts, *r_ij_* is the distance between the *i^th^* and *j^th^* residue, and Δ_*ij*_ = 1 for contacts in the tetramer. We decomposed *Q*(*t*) to *Q*(*t*)_*L*_ (contacts between residues with sequential separation between residues *i* and *j* less than or equal to 10, |*i* − *j*| ≤ 10), *Q*(*t*)_*LR*_ (fraction of global contacts, with |*i* − *j*| > 10), and *Q*(*t*)_*interface*_, which are interface contacts between residues *i* and *j* in different subunits.

Because each subunit is predominantly helical, the local contacts form rapidly. In contrast, both *Q*(*t*)_*L*_ and *Q*(*t*)_*LR*_ exhibit heterogeneous behavior. For a set of eleven trajectories (Fig. 7) we find that although the shapes of *Q*(*t*)_*LR*_ are similar, there is a large dispersion in their formation times. For example, the trajectory 11 (blue color in (Fig. 7) has not fully reached the equilibrium value in the dimer. Interestingly, there is a much greater diversity in the mechanism of formation of the interface residues. By comparing Fig. 7 and Fig. 8 we find that the approach of *Q*(*t*)_*interface*_ to the values in the assembled state varies greatly. In the orange trajectory in Fig. 8 one observes a rapid increase in *Q*(*t*)_*interface*_ from ≈ 0.2 to ≈ 0.8 in a single step. In contrast, we observe trapping in states with differing values of *Q*(*t*)_*interface*_ in the time interval between ≈ 100*μs* and ≈ 150*μs* in the purple trajectory. The dimer assembly is incomplete in a few trajectory on the 350 *μs* time scale. From these observations we conclude that trapping of the dimer in incorrect orientation is the primary cause for the slow assembly of the dimer.

**Figure 7:**
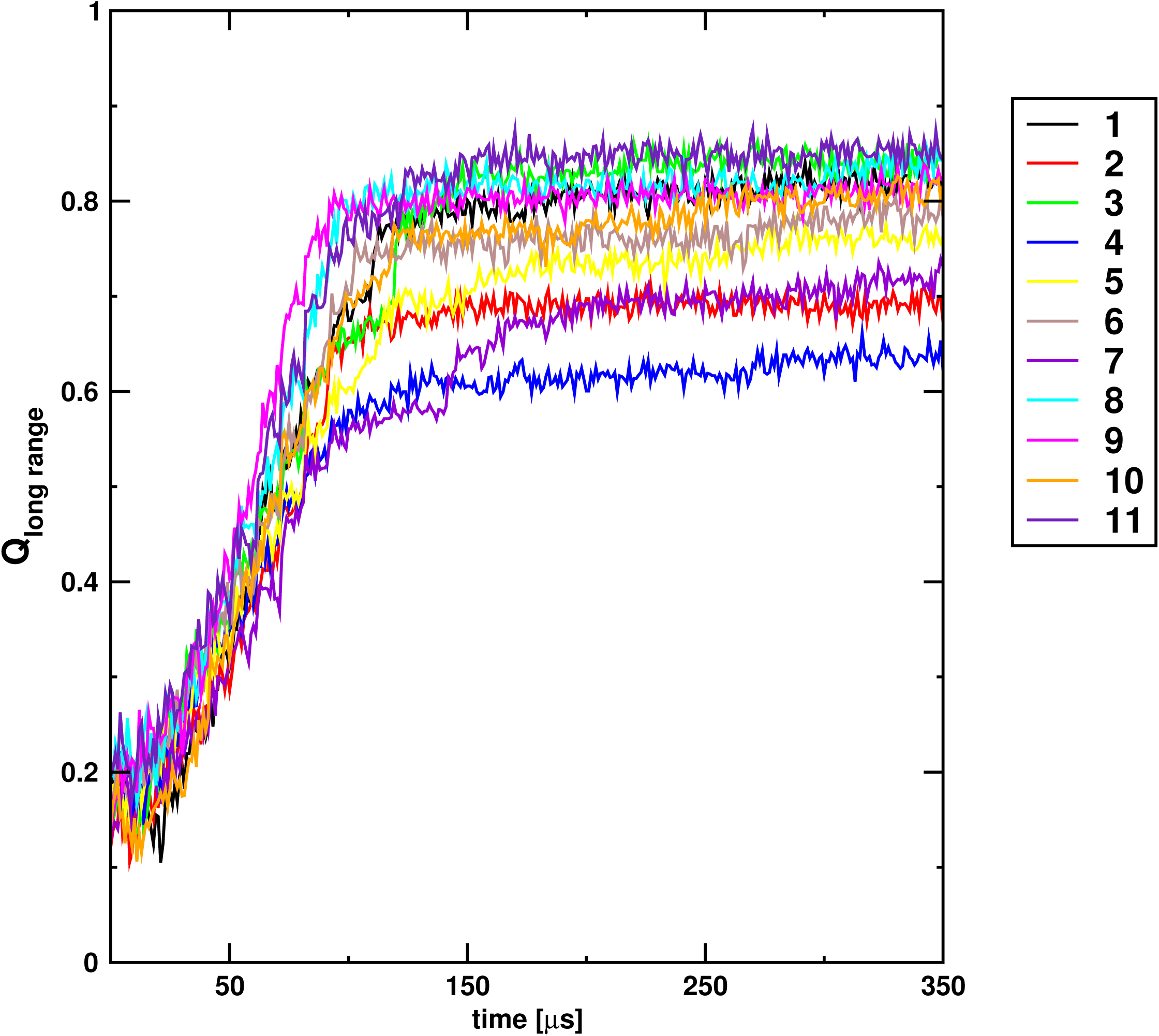
The fraction of native contacts as function of time calculated from a single refolding trajectory. *Q*(*t*)_*global*_ is plotted for 11 trajectories shown in black, red, green, blue, yellow, brown, violet, cyan, meganta, orange and indigo.

**Figure 8:**
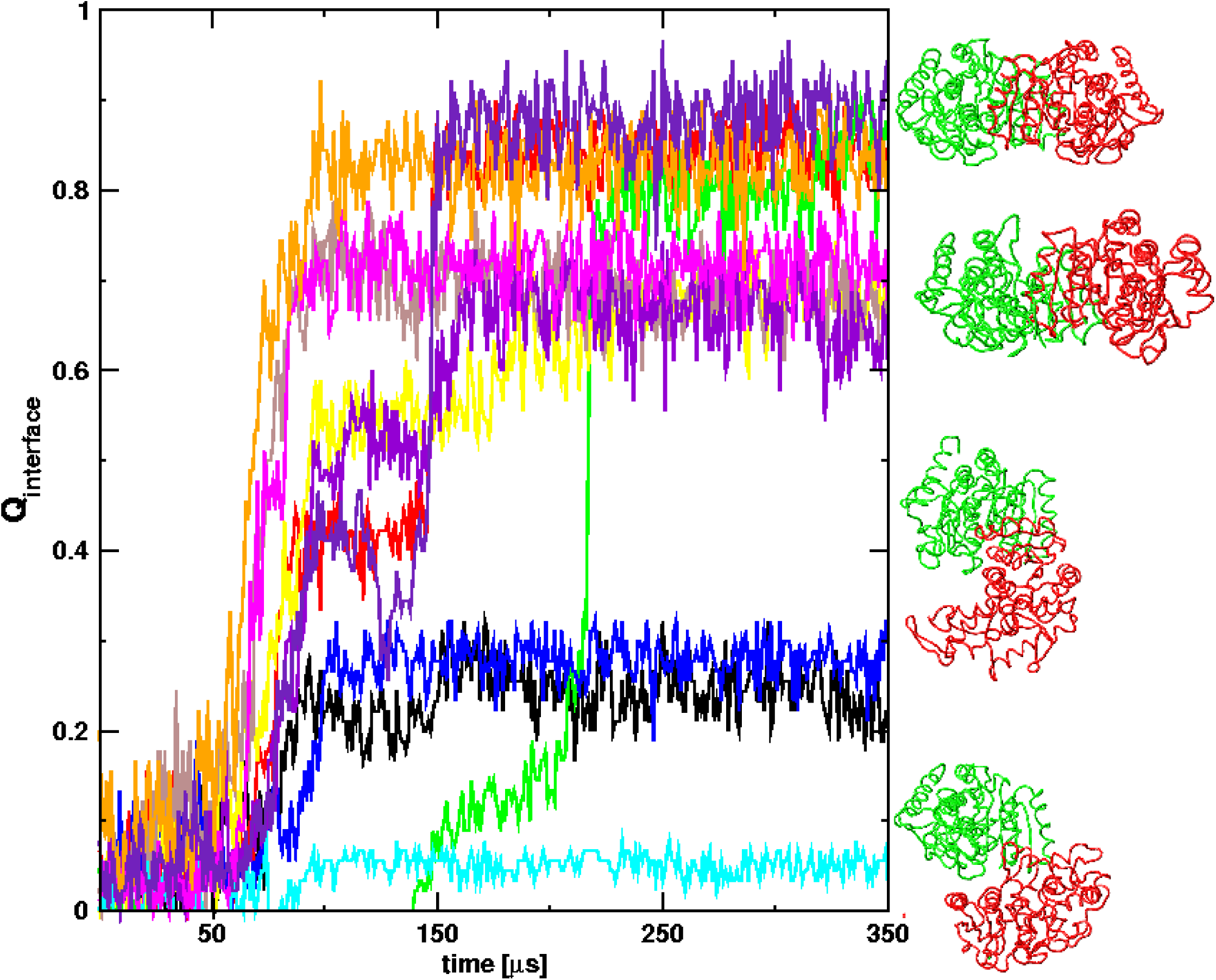
< *Q*(*t*)_*interface*_ > is plotted for the same 11 trajectories using the same color scheme as in Fig. 7. Structures of the trajectories in the end of the assembly process are shown from top to bottom. subunits 1 and 3 are colored in red and green, respectively.

Comparison of the dimer structures on the right of Fig. 8 shows that in all cases the individual subunits are formed but the extent of interface formation with correct orientation of the subunits varies substantially. Only in the purple and orange trajectories in Fig. 8 we find complete assembly of the dimers. The structures of the dimer at the right hand corner of Fig. 8 show that the interface is barely formed even though the individual are folded. Taken together, these results show that the rate limiting step is the formation of appropriate contacts between the interfaces that ensures proper orientational registry.

### Electrostatic interactions on the P-interface stabilizes the Q-interface

From the above results we conclude that the hydrophobic interactions on the Q-interface contributes to the stability of the dimer. In the predominant assembly pathway, prefolding of the subunits is required for dimer formation. However, the fully folded subunits could be arranged in an incorrect orientation, which results in pausing of assembly with intermediate *Q*(*t*)_*interface*_ values. The incorrectly oriented structures show that one monomer of the dimer is in the position where a P-interface related subunit in the tetramer LDH should be, which suggests that the P-interface interactions contribute to the correct orientation of Q-interface related subunits 1 and 3. An implication is that the symmetric tetramer structure should be more stable than the dimer structure. To prove this point, we performed additional simulations for the incorrectly oriented structures (cyan trajectory in Fig. 8) by including the P-interface related subunits (2 and 4). In Fig.9, *Q*(*t*)_*interface*_ in the cyan trajectory increases significantly from 0.05 to 0.7. The structures of the intermediate state of the assembly shows that the incorrectly oriented structures anneal in the tetramer assembly in contrast to the dimer formation in which the assembly is kinetically trapped due to energetic frustration. We surmise that the electrostatic interactions associated with the P-interface stabilizes the interaction on the Q-interface. Interestingly, the dynamics of allosteric transitions also involve the interplay between charged and hydrophobic in traction across the P and Q interfaces respectively, leading to the surprising conclusion that the network of residues controlling allosteric transitions and the assembly of LDH are nearly the same.

**Figure 9:**
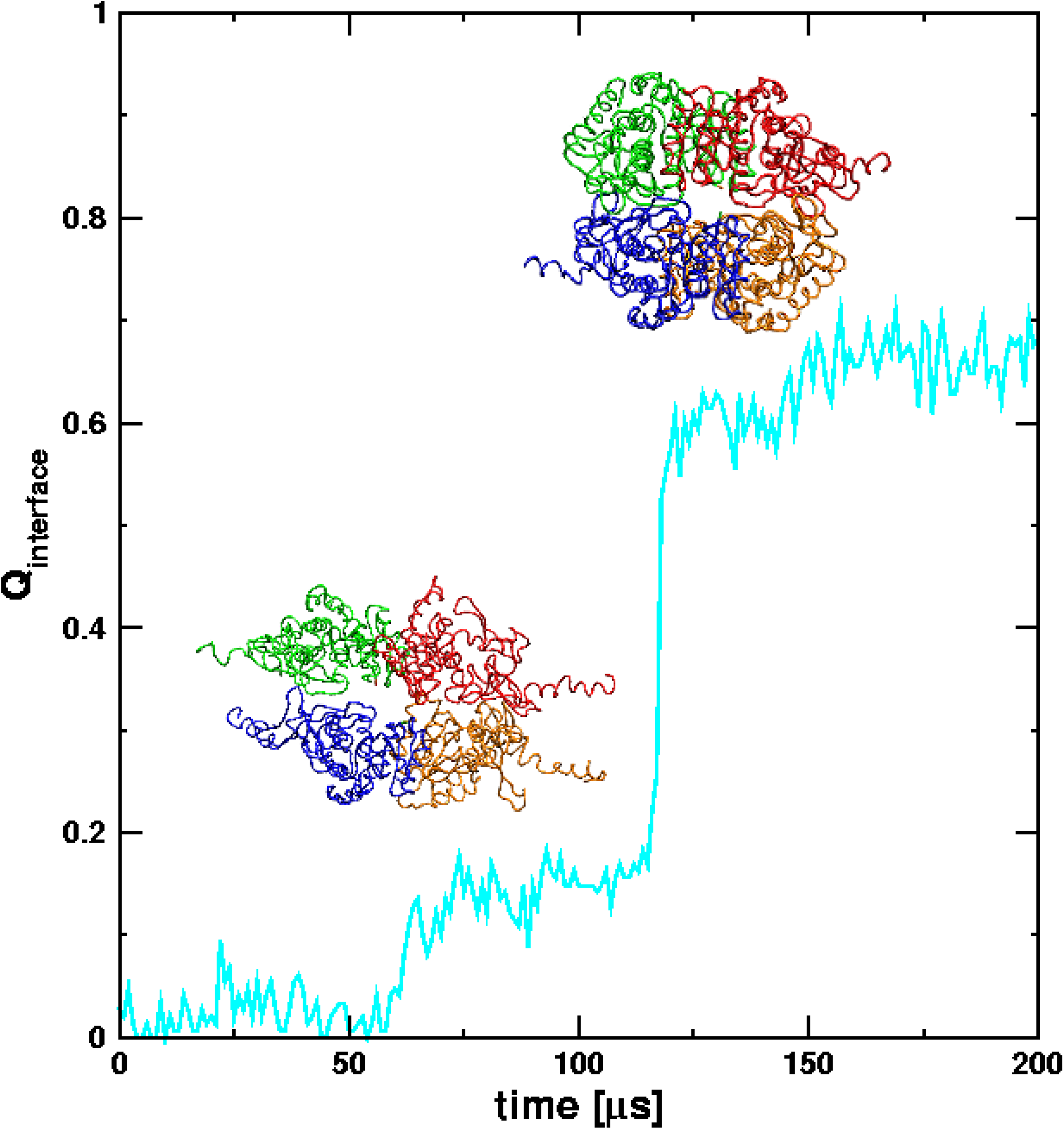
Time-dependent contact between interface residues, *Q*(*t*)_i*nterface*_, of the tetramer during the assembly process is plotted for the cyan trajectory. A snap shot of the metastable intermediate with *Q*(*t*)_*interface*_ ≈ 0.2 shows that the individual subunits have nearly adopted native-like structures but the overall assembly is far from complete. Assembly process is highly cooperative when the correct orientally registry is achieved, which in this case occurs at *t* ≈ 125*μs*. The subunits 1-4 are colored by red, orange, green and blue, respectively.

## Conclusions

The evolution of multi-domain allosteric proteins poses a number of questions such as what are the key interactions that drive transitions between distinct allosteric states, the mechanism of their assembly, and the relation between the two seemingly different processes. Using L-lactate dehydrogenase as a case study, we have established that many of the key residues in the allostery wiring diagram, identified using solely the fully assembled structures, that drive the dynamics of transitions between the T and R states, are localized at the interface between the subunits. Surprisingly, these residues also play an important role in the rate determining step in the assembly from unfolded structures. They ensure that the relative orientations of the subunits are correctly formed. The interface residues, which in the case of LDH are also involved in the allosteric transitions, might be evolutionarily conserved just as found for GroEL^26^. Our simulations also predict the temporal sequence changes in the secondary structural elements. We find that that the most dramatic secondary structural changes occur in neighborhood of the the active site.

It is unclear if the major conclusion that the residues in the AWD, predictable using structures alone using the SPM, not only drive the dynamics of allosteric transitions but also participate in the assembly in LDH, holds for other multi-domain allosteric enzymes. Our previous studies on GroEL^44^ and Dihydrofolate reductase^36^ have shown that the hotspot residues that are involved in the allosteric transitions coincide with those identified to transmit allosteric signals by the SPM. From the principle of parsimony, we surmise that the conclusions reached here for LDH might generally hold for other multiple-domain proteins. Combination of SOP simulations for the dynamics assembly and allosteric transitions and the use of SPM could be used to further test the generality of our conclusions, for multi-domain ring structures. The corresponding evolutionary requirements for linearly arranged subunits (Titin for example) may well be different, which we hope to investigate in the future.

## Acknowledgments

We are pleased to dedicate this paper to Bill Eaton, an extraordinary scientist, whose steadfast belief that solution of major problems in biology and even medicine requires physics and physical chemistry concepts has been a source of inspiration to the senior author for decades. This work was completed while JC was a graduate student at the Institute for Physical Sciences and Technology, University of Maryland. We acknowledge the National Science Foundation (CHE 16-36424) and the Collie-Welch Foundation (F-0019) for supporting this work.

---

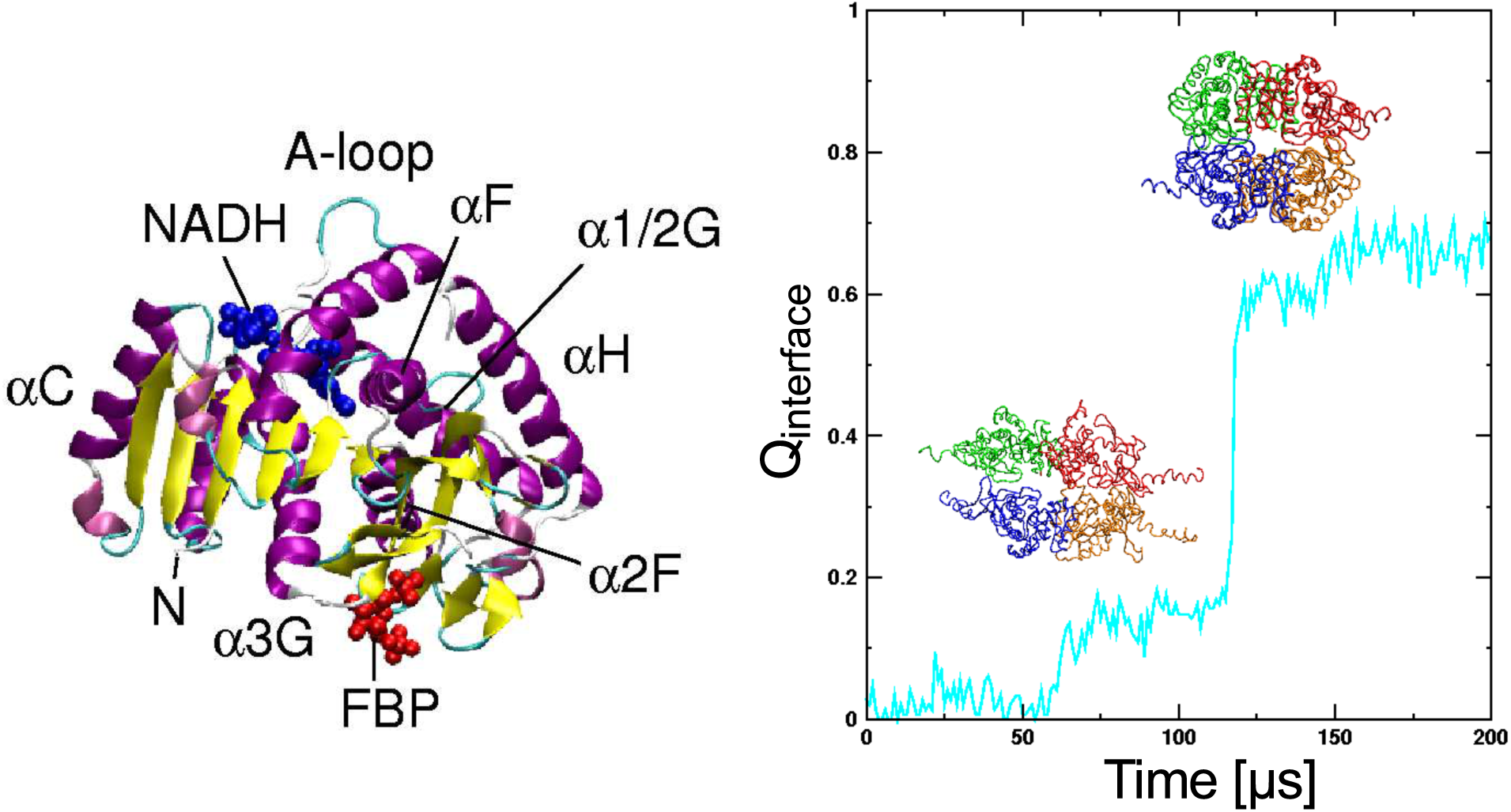
Table of Contents Graphics

